# Generative Artificial Intelligence for Secure and Scalable Multimodal Eye Movement Datasets

**DOI:** 10.1101/2025.04.21.649786

**Authors:** Aimon Rahman, Preetham Bachina, Vishal Patel, Kemar E. Green

**Affiliations:** Johns Hopkins University, Department of Electrical and Computer Engineering, Baltimore, MD, USA; Johns Hopkins University, School of Medicine, Baltimore, MD, USA; Johns Hopkins University, Department of Neurology, Baltimore, MD, USA

## Abstract

Eye movements, such as nystagmus, saccades, and smooth pursuit, provide valuable information about neurological function but have limited publicly accessible datasets due to patient privacy concerns. To address this, we leverage generative AI to create realistic videos of artificial eye movement, eliminating the need for real patient data. These synthetic datasets have shown performance comparable to actual patient data in clinical tasks. Our generated videos will be openly shared, facilitating broader research and advancement in neurologic and neuro-ophthalmic AI applications.

## 1 Introduction

Eye movement disorders can be markers of neurologic disease. [6]. They develop from disruptions within brain-eye-ear circuits, which are ubiquitous throughout the brain. Recent research utilizing machine learning algorithms has attempted to detect and analyze eye movements from video-oculography (VOG) data (video and waveform) [13, 12, 10, 11, 14, 21, 24, 9]. However, these studies often use small, limited datasets with few variations and are generally not publicly available due to privacy concerns[8][7][25], which hinders further research. A common approach to address the scarcity of public datasets in the medical field is the generation of synthetic datasets using real patient data [2,], a method that has been proven effective in various downstream tasks. However, generating synthetic data from real-world datasets presents its own challenges. For example, generative models can sometimes replicate training data too closely, undermining the purpose of creating synthetic datasets. [18]. Lastly, creating clinically relevant eye movement datasets requires generating long-form videos that accurately integrate oculographs (pupil position waveforms) with physiological eye movements, adding complexity to the task.

**Figure 1:**
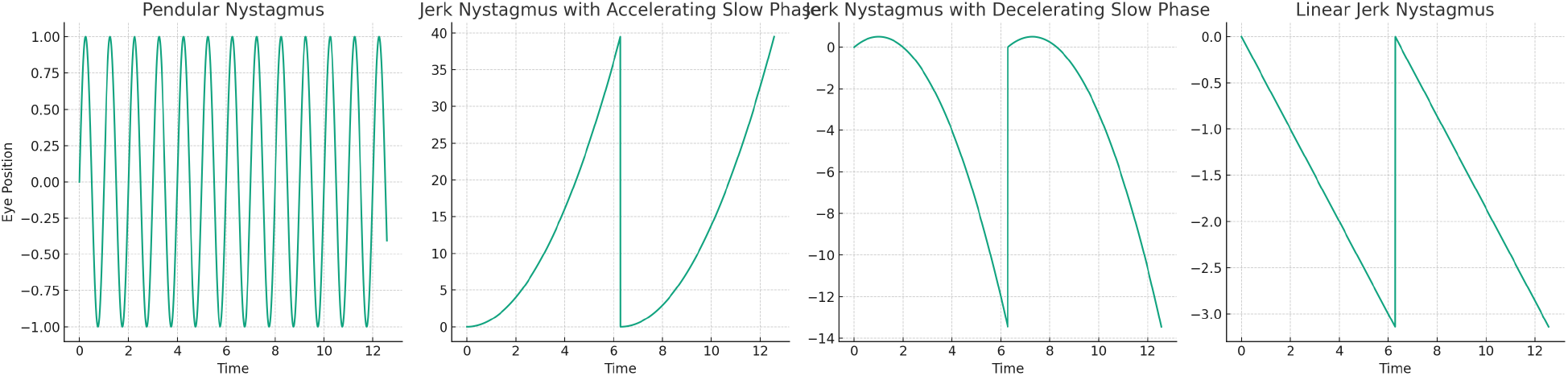
Waveform morphologies of various nystagmus types. Nystagmus can be categorized into two main types based on the pattern of eye movements: jerk nystagmus, which consists of a slow phase followed by a quick phase, and pendular nystagmus, characterized by movements that are slow in both directions. These patterns reflect the underlying issues with the neural pathways responsible for eye stability. Additionally, jerk nystagmus can be subdivided into three categories based on the speed of the slow phase: constant or linear velocity, decelerating velocity, and accelerating velocity.

In this paper, we address the challenge of creating long-form nystagmus videos without compromising patient data by developing our system. Our key contributions are as follows.

- We successfully generate synthetic videos of nystagmus encompassing a range of types of nystagmus, overcoming the limitation of being restricted to a single or specific type.
- Our method avoids the use of real patient data, thus preserving patient privacy. This approach is especially valuable in scenarios where data availability is limited. We validate our dataset using real patient data for comparison and engage experts for further validation.
- We made the synthetic data set publicly available, in order to improve the research and understanding of nystagmus in the broader scientific community.

## 2 Methods

### 2.1 Preliminaries

#### Nystagmus Waveforms

In the following section, we present equations representing pupil location over time for different types of nystagmus, as illustrated in the graphs in Figure **??**. The variables in these equations are consistent across all types: *A* represents the amplitude, indicating the maximum extent of eye movement; *ω* is the angular frequency, determining the speed of oscillation; *ϕ* is the phase, accounting for any initial positional offset; and *t* denotes time.

1. *<u>Pendular Nystagmus</u>*: Characterized by rhythmic, oscillatory eye movements, similar to the motion of a pendulum. It is described by the equation: *P* (*t*) = *A ×* sin(*ωt* + *ϕ*).
2. *<u>Jerk Nystagmus with Accelerating Slow Phase</u>*: This type involves eye movements that slowly accelerate in one direction and then quickly jerk back. The equation representing this motion is: *J*_*a*_(*t*) = *A ×* (*t*^2^ + *ϕ*).
3. *<u>Jerk Nystagmus with Decelerating Slow Phase</u>*: In this variation, the eye slowly moves in one direction with a gradually decreasing speed before jerking back. This is characterized by the equation: *J*_*d*_(*t*) = *A ×* (*t −* 0.5 *× t*^2^).
4. *<u>Linear Jerk Nystagmus</u>*: This type features linear, steady eye movements in one direction followed by a rapid jerk in the opposite direction. It is described by: *J*_*l*_(*t*) = *A ×* (*−*0.5 *× t*).

### 2.2 Modeling Nystagmus Pupil Movement

In the process of modeling videos with the pupil kinetic patterns of nystagmus, the initial step involves choosing a specific nystagmus waveform equation, symbolized as *f* (*t*; *θ*). Here, *t* signifies time, and *θ* encompasses parameters such as amplitude, frequency, phase, among others, which are essential for shaping the waveform. Upon selecting the waveform equation, a binary mask video *V* (*t, x, y*) is crafted to depict the pupil’s trajectory over time. This is accomplished by calculating the pupil’s coordinates (*x*(*t*), *y*(*t*)) for each time instance *t*, based on the waveform equation. Subsequently, the binary mask video is updated to indicate the pupil’s position, with *V* (*t, x*(*t*), *y*(*t*)) = 1 marking the pupil’s location, and *V* (*t, x* ≠ *x*(*t*), *y* ≠ *y*(*t*)) = 0 for all other points, effectively distinguishing the pupil from the remainder of the video frame. If we were to represent this as a continuous process, the creation of the binary mask video could be expressed as:

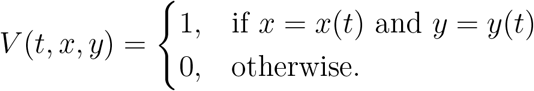

This method outlines a systematic approach to dynamically map the pupil’s movements across time, utilizing the nystagmus waveform equation to precisely replicate this ocular phenomenon.

Since we are not utilizing any patient dataset with nystagmus eye movement, we generate pupil movements. The procedure begins by selecting a random equation that corresponds to a type of nystagmus waveform. We then adjust parameters such as the amplitude and other variables to shape the specific waveform. Following this, we generate a binary mask video. In this video, the pupil’s location changes over time in accordance with the selected equation, thereby replicating the distinctive waveform of the nystagmus type in question. This approach results in a series of masked videos, each representing a different kind of nystagmus waveform. We employ these masked videos as a basis for generating synthetic nystagmus videos. Using masked videos, which depict varying pupil locations over time according to specific nystagmus waveforms, we can condition the synthetic video generation process.

### 2.3 Eye Movement Video Generation

#### 2.3.1 Network Architecture

The core of this model is the latent video diffusion mechanism, essential for the conversion of noise to visual representations [15, 22]. This process is orchestrated by a U-Net structure, labeled *ε*_*θ*_. A key element in this architecture is the VQGAN, which serves to bridge the visual and latent domains. Given a training video *v*^*gt*^ in RGB format, it is first encoded by the VQGAN encoder, *ε*, mathematically represented as

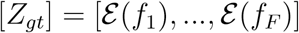

where *v*^*gt*^ = [*f*_1_, …*f*_*F*_] symbolizes the frames of the training video. The U-Net, denoted as *ε*_*θ*_, operates in the latent space, iteratively refining the video’s representation through a fixed number of *T* steps. During inference, the model’s primary goal is to predict noise as

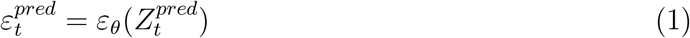

with 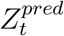 being the latent variable at the *t*-th step. The predicted denoised latent *Z*^*pred*^ is eventually decoded through the VQGAN decoder 𝒟 and projected back into the pixel domain. The efficacy of the denoising UNet is evaluated using the loss function

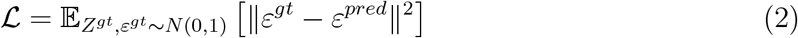

where *ε*^*pred*^ and *ε*^*gt*^ signify the predicted and ground-truth noise, respectively. The U-Net architecture encompasses various segments, including initial, down-sampling, spatio-temporal, and up-sampling blocks. The spatio-temporal block, crucial for video generation, captures spatial and temporal dynamics in eye movement videos [22]. Process individual frames, employing spatiotemporal attention within a transformer architecture [1], to discern spatial details and their sequential progression. In the spatial attention block of the multi-head attention layer, we utilize the text embedding of the prompt as both the key and the value.

### 2.4 Training Process

#### Learning to Generate Eye Movement Videos

At this phase, we leverage a publicly accessible eye dataset LPW [20] to train the video diffusion model, using the prompt “Eye with Normal Pupil Movements” for each video. The training process is carried out in 10,000 steps, with a learning rate set at 5 *×* 10^*−*5^.

#### Learning the Nystagmus Motion

In this phase, we exclusively fine-tune the temporal layers on Nystagmus pupil videos obtained from Section 2.2, allowing the model to learn nystagmus movement without changing the spatial information. This pre-trained model is already adept at creating eye videos. Thus, by training the temporal block, we enhance the model’s ability to learn eye movements of the nystagmus from the dataset, while avoiding the replication of the specific content of the dataset. This stage is trained for 1000 steps, with the same learning rate as before. In the initial setup, we train a model expert in generating eye movement videos, abstracting any pathological deviations such as nystagmus.

**Figure 2:**
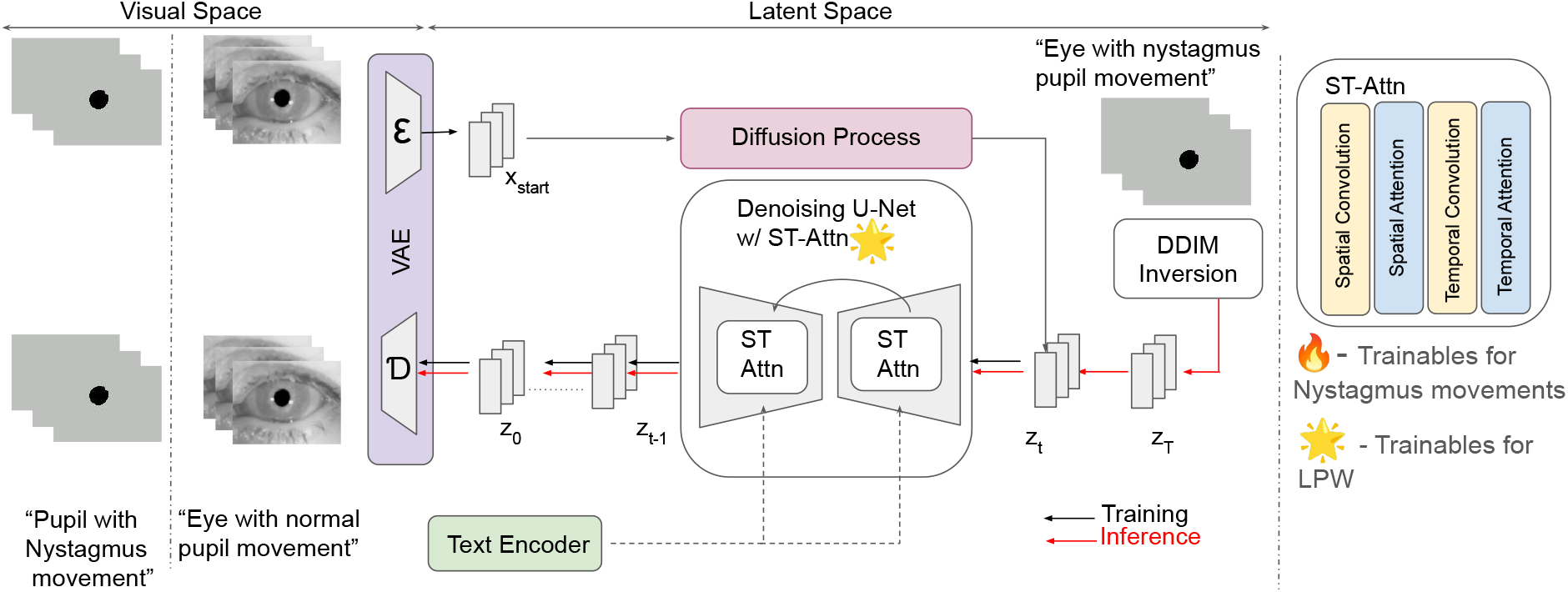
The figure illustrates the architecture and workflow of the proposed method, beginning with training the entire model on the LPW datasets to capture natural eye movements and surrounding orbital features. Next, we freeze all layers except for the temporal ones, which are then trained to detect the motions characteristic of nystagmus from pupil videos. The inference process utilizes the pupil videos and textual prompts to produce nystagmus videos.

### 2.5 Inference: Conditional Nystagmus Video Generation

In the process of video generation, especially for creating eye-movement videos, starting from pure noise yields impressive results. However, using only noise as a basis offers limited control over the specific waveform patterns in the generated video. To enhance control, we incorporate movement cues derived from video waveforms. These cues are obtained from masked videos generated using waveform equations during the inference stage. Specifically, we introduce noise to the latent representations of these masked videos, denoted *v*_*m*_, and then apply a denoising process. This step is performed using the Denoising Diffusion Implicit Model (DDIM) [19, 23] inversion technique. The added noise in the latents forms the basis for DDIM sampling, which is guided by the waveform characteristics. As a result, the final output video, represented as *V*, is formulated as follows:

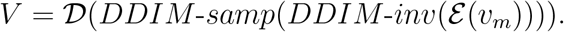

This approach allows for the creation of more controlled and specific eye movement patterns in the generated videos. [**?**]

### 2.6 Network Architecture

We propose the Shadow and Surface Segmentation Network (SSNet) for simultaneous bone surface and shadow segmentation from US images which is illustrated in Figure **??**. SSNet is composed of a shared LeViT-based encoder to extract global and long-range spatial features and two CNN-based decoders with a cross-task feature transfer block to leverage mutual information between the two tasks.

**Figure 3:**
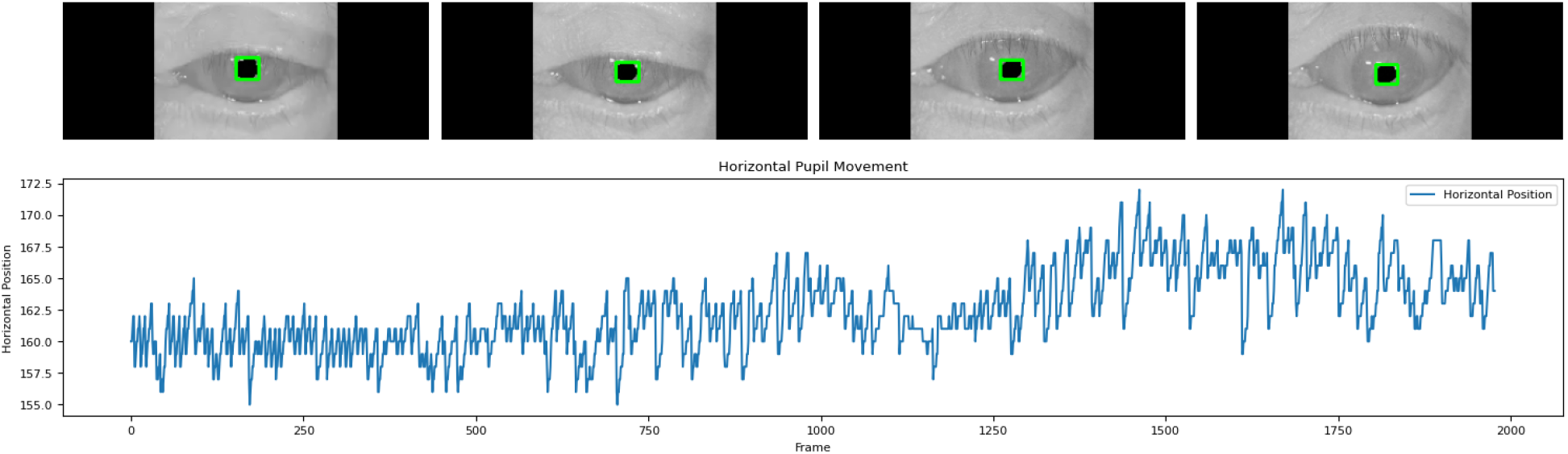
Nystagmus pose estimation

## 3 Experiments and Results

### 3.1 Implementation Details

In our approach, we utilize a video generation model that has been pre-trained, as outlined in [22]. This model is founded on the principles of Latent Diffusion Models, which are elaborated in [15], and it employs pretrained weights that are available to the public. We process the input video by extracting 256 evenly distributed frames, ensuring that each frame has a resolution of 256×256. Following this, we proceed to fine-tune our models by employing a method designed for learning motion. The entire training procedure is conducted with a batch size of 1. For the phase of inference, the DDIM sampler [19] is utilized, and we also incorporate classifier-free guidance [3] into our experimental setup. On average, generating 60 frames with an NVIDIA A100 GPU takes approximately 1 minute.

### 3.2 Datasets

#### Public Dataset

To train the model we utilize the Labelled Pupils in the Wild (LPW) [20] dataset that introduces a unique collection of 66 high-quality, high-speed videos focusing on the eye region, designed for the advancement and assessment of pupil detection algorithms. Recorded at approximately 95 FPS with a cutting-edge dark-pupil head-mounted eye tracker, these videos were captured from 22 participants across various everyday settings. The LPW dataset encompasses individuals from multiple ethnic backgrounds and features a wide range of indoor and outdoor lighting conditions, in addition to natural variations in gaze directions.

#### Private Dataset

In the validation study, we examined a cohort of 20 patients. Each participant contributed videos capturing movements of both the left and right eyes, with durations exceeding a minute in some cases. The data collection strategy is based on previously described methods [5].

#### Baselines

Given the absence of direct methodologies for inducing nystagmus in normal eye videos, we have identified VideoControlNet [4] as a foundational comparative method. This framework leverages a video generation network that operates under the guidance of a mask to produce subsequent video. We train a VideoControllNet [4] with our natural eye dataset for comparison. We utilized 60 videos labeled as nystagmus and 70 videos categorized under the normal class to train a ResNet-50 classifier aimed at disease identification. For testing purposes, we reserved another set comprising 54 videos of nystagmus and 43 of the Normal class from the private dataset.

**Figure 4:**
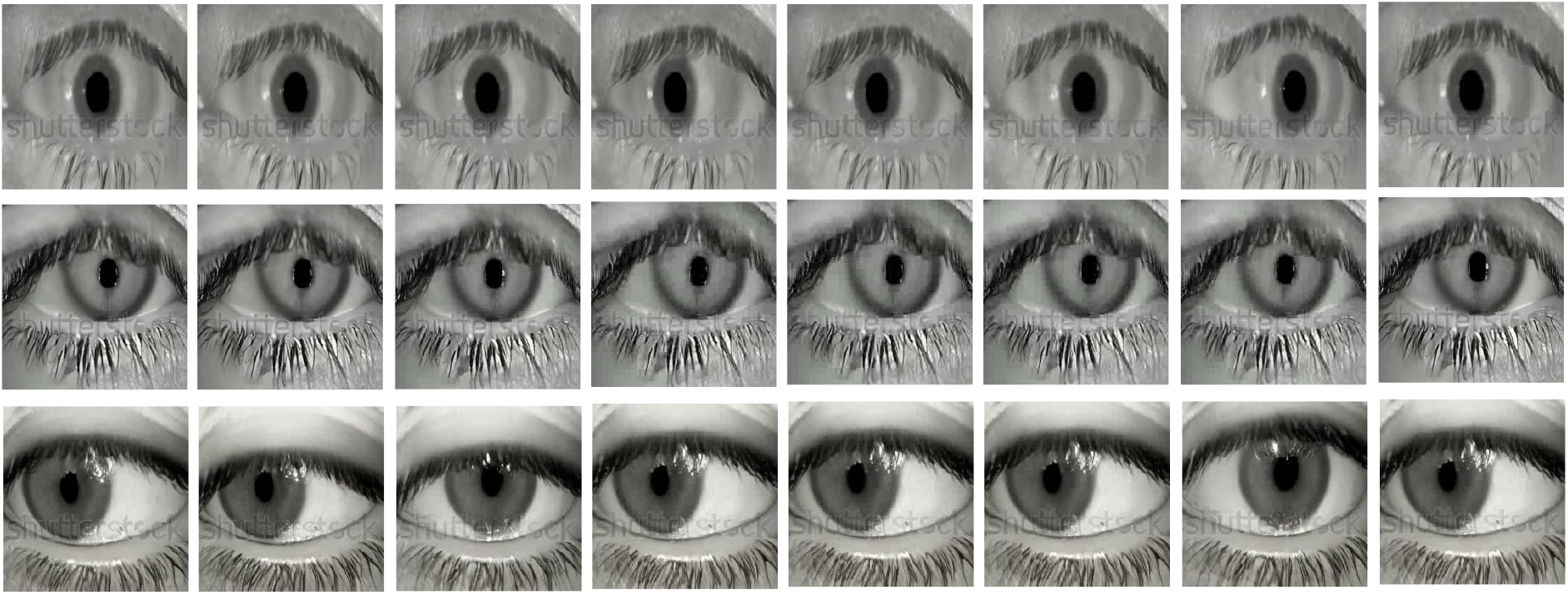
Sample frames extracted from the synthesized nystagmus video sequences

### 3.3 Quantitative and Qualitative Analysis

For the quantitative evaluation, we used the Fréchet Video Distance (FVD) as our assessment tool, achieving a score of 295. To further validate generated videos, we proceeded to train a classifier leveraging the synthetic dataset using both ControlNet and our approach. These findings are detailed in Table **??**. The videos generated by ControlNet [4] show more artifacts and appear more mechanical, so the FVD remains higher. In comparison, our method demonstrates a lower FVD score as well as higher downstream accuracy. We also report the downstream performance of the full dataset as a reference. Figure **??** displays frames extracted from the generated videos, showcasing the network’s ability to create high-quality videos.

**Table 1:** Quantitative results of the generated videos in terms of FVD and their impact on downstream performance. Real dataset indicates a classifier trained on the full dataset and tested on the test set.

| Approach | FVD (↓) | Downstream Accuracy (↑) |
| --- | --- | --- |
| ControlNet [4] | 971 | 74.32 |
| Real Dataset | — | 84.12 |
| Ours | 295 | 77.32 |

### 3.4 Ablation Studies

In this section, we evaluate the significance of each component within our network, particularly during the inference process, with the findings presented in Table 2. First, we employ only the DDIM inversion technique on masks to produce nystagmus videos, independent of any prompt input. Although this method can occasionally produce nystagmus videos, it often misses the waveform influence, leading to diminished performance in downstream tasks. Next, we experiment with using prompts alone to direct the generation process, essentially denoising pure Gaussian noise to create the eye video. This approach results in suboptimal downstream accuracy, as certain videos do not exhibit accurate movement patterns. These experiments demonstrate the necessity of integrating prompt guidance and DDIM inversion for effective video generation.

**Table 2:**
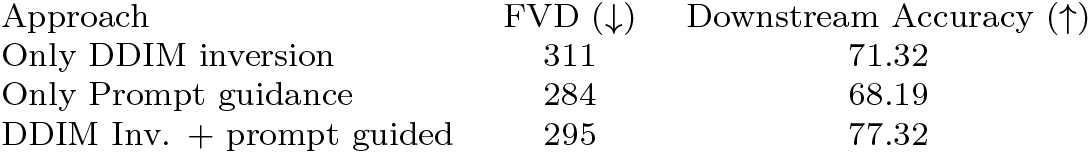
Ablation Studies.

### 3.5 Comparison with Motion-Controlled Video Generation Techniques

A common method in motion-controlled video generation involves the utilization of VideoControllNet. This technique allows for the extraction of the pupil, enabling the training of the Video Diffusion Model with the movement of the pupil as a conditional factor. Specifically, the Nystagmus pupil movements are incorporated during the inference phase. Employing this methodology, we developed a model that produces videos with a mechanical essence, mirroring the entire waveforms without any natural eye movement patterns. The videos generated using this approach yielded an FVD score of 971.

### 3.6 Impact on Downstream Task

In this section, we assess the utility of the nystagmus videos generated for a downstream task. Initially, we used synthetic videos to train a classifier specifically designed for nystagmus detection. Subsequently, we test this classifier using a real-world dataset comprising patients diagnosed with nystagmus. As evidenced in Table X, the classifier shows a high precision in identifying patients with nystagmus. This result substantiates the effectiveness and reliability of the synthetic dataset in practical applications.

### 3.7 Training data Replication

A significant challenge in synthetic data generation is the risk of sample replication [18], which can lead to the memorization of training data and the potential leakage of patient information. Despite using a public eye dataset for this project, we diligently examined the possibility of such a replication. We report the highest average SSCD scores (Self-Supervised Copy Detection) comparing the training data set and the generated data, which are 46. 7%, indicating minimal similarity, which confirms a reduced risk of data replication.

### 3.8 Limitation and Ethical Impact

A notable limitation of the model is its occasional production of videos with unrealistic features, such as eyes that possess two pupils. Similarly, we have not modeled the nystagmus based on the expected physiological changes that occur in its amplitude and velocity during normal or pathologic voluntary or involuntary eye movements[6][16][17]. This requires manual review to remove such unrealistic videos. Given the stochastic nature of the sampling process, it is also prone to introduce artifacts. In addition, video generation technologies, particularly sophisticated video diffusion models, pose ethical and social challenges. These challenges encompass the potential for generating misleading content, which could contribute to misinformation and infringe on privacy rights; the risk of biases in AI-generated content, which could result in discriminatory or unfair outcomes; and concerns regarding the impact on intellectual property rights. It is imperative for both developers and users of these technologies to recognize these risks and commit to their responsible utilization.

#### Implementation Details

SSNet is trained using a batch size of 32. For training both branches, a two-step training phase is adapted. Each of these steps are trained until convergence. The weights and bias of the network are optimized using the Adam optimizer with a learning rate of 10^*−*4^. All US scans and their corresponding masks are resized to 224 *×* 224 pixels and rescaled between 0 and 1. All transformer blocks in the LeViT architecture were pre-trained on ImageNet-1k. The overall loss function we use to train the multi-task network is,

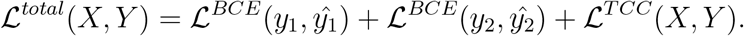

Binary-cross entropy loss has been used between the prediction and the ground truth, which is expressed as,

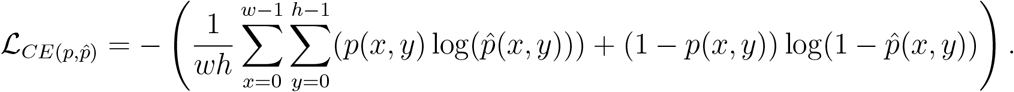

## 4 Conclusion and Future Direction

In this research, we present a novel method for generating synthetic nystagmus video-oculography (VOG) data without relying on patient-specific information, instead leveraging a publicly accessible dataset. We developed video diffusion models capable of producing various types of nystagmus by integrating both waveform and prompt conditioning. Utilizing a large-scale public dataset ensures patient privacy and mitigates concerns regarding data duplication through pre-trained generative models. We plan to share these synthetic videos and corresponding waveforms openly with the research community for further validation. Future work will focus on the comprehensive validation of synthetic videos and the creation of additional nystagmus variants that incorporate realistic noise patterns. Our approach represents an important step forward in the deep learning-based detection of eye movements associated with neurologic and ophthalmologic diseases.

## Acknowledgements

We gratefully acknowledge funding support from the Johns Hopkins Institute for Data-Intensive Engineering and Science (IDIES) and the Johns Hopkins University (JHU) Discovery Award. Computational resources were provided through the Advanced Research Computing at Hopkins (ARCH). We also extend our appreciation to Johns Hopkins University for its overall institutional support and resources that facilitated this research.

## Notes

### Competing Interest Statement

The authors have declared no competing interest.

